# Synergic homology directed recombination by PRDM9 meiotic factor

**DOI:** 10.1101/2022.12.05.519167

**Authors:** Marta Sanvicente-García, Lourdes Gonzalez-Bermudez, Isabel Turpín, Laura Batlle, Sandra Acosta, Marc Güell, Avencia Sanchez-Mejias

## Abstract

Genome editing requires precision to broadly move on to industrial and clinical applications. For this reason, homologous directed repair (HDR) is one of the preferred methods for small edits, other than knock-outs. However, HDR has low efficiency. Current investigations to enhance HDR have mainly gone in the direction of finding non-homologous end joining (NHEJ) inhibitors. NHEJ is crucial for cellular integrity, then the inhibition of this pathway is detrimental for the correct survival of living entities. In other studies, a second opportunity is given to HDR by targeting the byproducts of NHEJ, using an extra gRNA. In this study, we propose the use of a meiotic factor, PRDM9, to directly enhance homology recombination. Through the exploration of combinatorial factors and donor design, we have established an optimized protocol for HDR. PRDM9-Cas9 fusion combined with CtIP improves HDR/NHEJ ratio. In addition, we have validated this combinatorial approach for small edits through a traffic light reporter system, as well as for longer edits with a split-GFP reporter system.

## INTRODUCTION

Small genome edits using homology directed repair (HDR) can be performed by introducing a gRNA that targets a position near to the target locus, in conjunction with Cas9, and a donor template DNA (Heyer, Ehmsen, and Liu 2010). The insertion of precise edits to the human genome is limited by the relatively low efficiency of HDR compared with the often higher efficiency competing DNA repair pathway of non-homologous end-joining (NHEJ) (Heyer, Ehmsen, and Liu 2010).

Researchers have explored numerous approaches to potentiate HDR. Inhibition of NHEJ by targeting multiple components, such as KU70, KU80 or DNA ligase IV, can increase up to eightfold HDR (Chu et al. 2015). Small molecule DNA-PK inhibitors enable similar increases in HDR (Robert et al. 2015). Alternative approaches include potentiating HDR components such as RS-1, which enables increases of up to 5-fold (Song et al. 2016), or also small molecule screening led to the discovery of HDR potentiators showing up to 9-fold increase in pluripotent stem cells (Yu et al. 2015). There are other molecules that enhance HDR by inducing cell cycle arrest at G2/M phase like histone deacetylase inhibitors (Li et al. 2020) by the increase of DNA accessibility (Liu et al. 2020). The discovery of 53BP1 inhibitors, such as DP308 could also increase CRISPR-mediated genome editing precision (Sun et al. 2021). In other studies, the importance of the colocalization of the donor and the Cas9 have been highlighted (Li et al. 2021). The use of a Cas9 transcription factor fusion and adding THAP11-specific DNA binding motifs to both ends of the double-stranded donor DNA promotes HDR. Up to 6-fold increases of HDR is shown through the combinational use of the transcription factor fused to Cas9 and valnemulin, a small-molecule HDR enhancer (Li et al. 2021). XL413, an inhibitor of CDC7 has also been shown to increase HDR efficiency in primary T cells (Wienert et al. 2020), as well as in iPSCs (Maurissen and Woltjen 2020). Even anti-CRISPR proteins fused to CDT1, which is degraded in S and G2, have been used to promote editing in the cell cycle when HDR is dominant (Matsumoto, Tamamura, and Nomura 2020). In this direction, the design of the template DNA has also been exploited to maximize targeting. Targeting the strand that is released first after cleavage, HDR can increase targeting 5-fold (Richardson et al. 2016). Last efforts in enhancing HDR have been done by targeting NHEJ byproducts using a second gRNA to target the most frequent indels (Bodai et al. 2022) (Möller et al. 2022). Even giving a second chance to HDR, through gRNAs targeting NHEJ products, can be efficient, the usage of extra gRNA can be detrimental when accurate and traceable results are required, such as in clinical CRISPR applications and environmental or industrial uses. Since the very early use of CRISPR-Cas9 for mammalian cell genome editing, concerns for the off-target effects have been raised (Fu et al. 2013). Moreover, new sequencing techniques have enabled the discovery of previously unseen off-target patterns (Höijer et al. 2020), and there are cases where the use of multiple gRNAs have led to translocations (Samuelson et al. 2021). Then, the probability of unspecific targets can be increased each time that a new gRNA is added to the genome editing design. Anyway, in cases where off-targets from the extra gRNA are highly improbable, this can be a complementary strategy to the use of factors enhancing HDR or inhibiting NHEJ.

Human cells undergo natural efficient recombination during meiosis after double stranded DNA breaks. This process is efficient and safe. Understanding this process can help us to harness the natural factors to engineer safer and more efficiency editing tools. Multiple factors have been described to influence meiotic recombination. PRDM9 is a natural fusion zinc finger – chromatin remodeler that kicks off recombination hotspots in meiosis (Baudat et al. 2010). Recombination efficiency depends on having PRDM9 bound to the template at the recombination site, and having the double stranded cleavage performed by spo11 (Hinch et al. 2019). It could be possible to emulate this natural system to increase the HDR outcome in the context of genome editing with Cas9 and PRDM9 fusions, however this has not been explored in the past. Interestingly, dead Cas9 has been previously fused to the catalytic domain of PRDM9 to achieve transcriptional activation by means of the methylation activity to H3K4 to create transcriptional activation and prevent silencing (Cano-Rodriguez et al. 2016).

In the present study we wanted to explore the potential of PRDM9 to increase HDR in combination with Cas9, and other factors known to increase HDR efficiency for genome editing applications. We have demonstrated the capacity of PRDM9 fused to Cas9 of favoring HDR over NHEJ after a DSB, both in the context of small DNA edits and site specific integration of several kilobases long donor DNAs. We further explored the synergy of PDRM9 mediated HDR enhancement with other effectors known to favor HDR, including template design, in human fibroblast and iPSC cells. We have demonstrated the potential of the chromatin remodeler PRDM9 to favor HDR, opening new possibilities for the design of efficient genome editing tools.

## RESULTS

### PRDM9 as an HDR promoting factor validated with traffic light reporter (TLR) and split-GFP reporter system

We first constructed a C-terminal fusion of *S. pyogenes* Cas9 and PDRM9 and tested recombination efficiency with a split GFP reporter system previously described (Fig. 1A; (Pallarès-Masmitjà et al. 2021)), both as episomal DNA (Fig. 1B) and in cells containing the split GFP reporter in the genome (Fig. 1C). We observed an increase of HDR of 40% (Fig. 1B and C, p-value=0.04 and 0.03).

**Figure 1.**
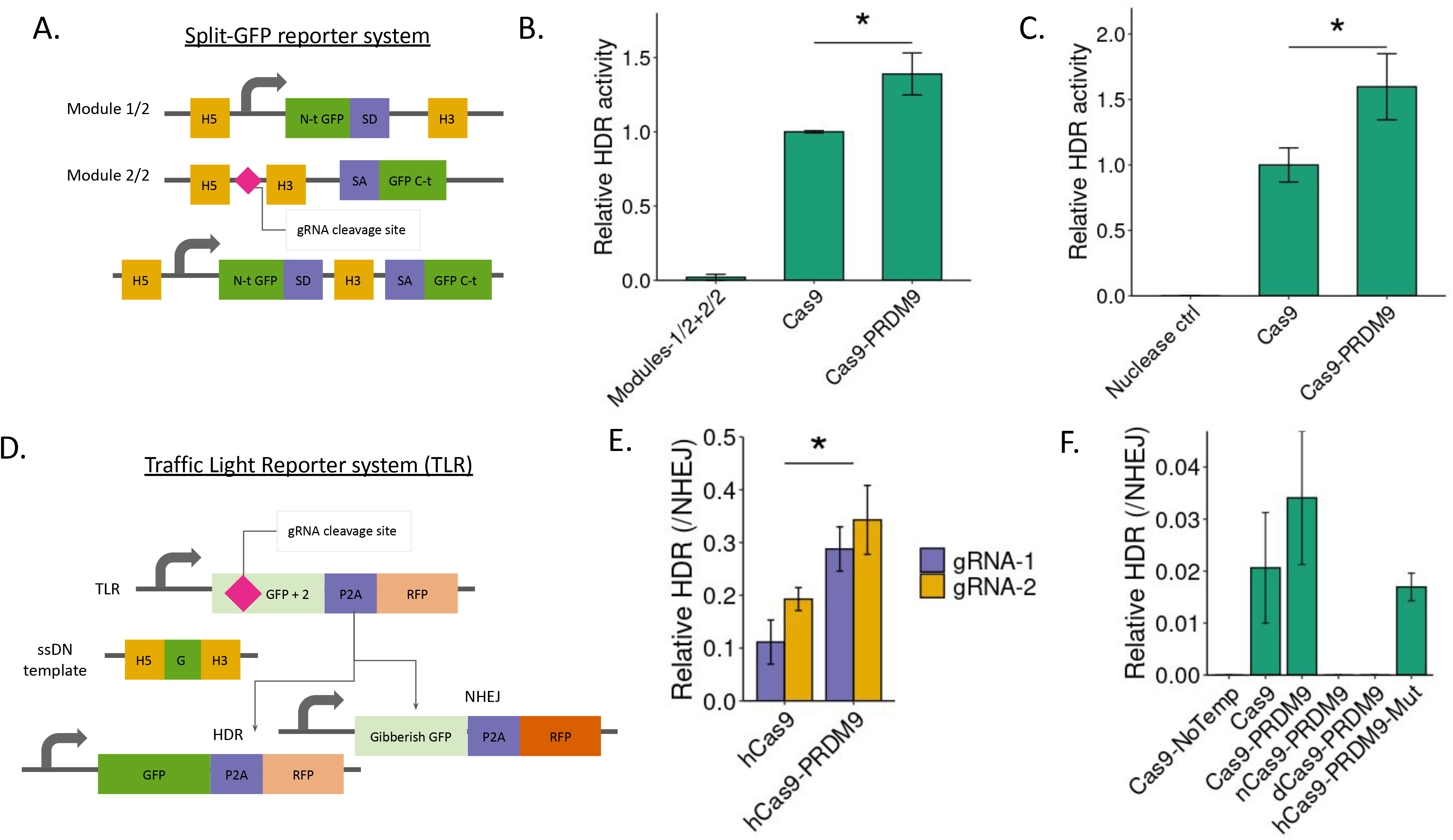
Split-GFP and traffic light reporter systems to validate PRDM9 as an HDR promoting factor. A) Scheme of split GFP reporter system. H5 stands for 5’ homologous recombination sequence, H3 for 3’ homologous recombination sequence, SA splicing acceptor, SD splicing donor. B) HDR activity measured using the split-GFP reporter when acceptor split GFP construct is provided as an episomal plasmid (p=0.042); or C) integrated in the genome (p=0.033). D) Scheme of the traffic light reporter system to measure HDR and NHEJ efficiencies. E) HDR relative to NHEJ measurement comparing either hCas9 alone or in fusion with PRDM9, F) or PRDM9 fused with nuclease, nickase and dead variants of Cas9 as well as catalytic mutant PRDM9.

An alternative traffic light reporter system was used to estimate the increase in HDR compared with NHEJ (Fig. 1D). We generated a stable cell line with a single copy of the TLR system in Hek293T cells. Although HDR absolute efficiency did not change significantly in the conditions tested, a significant decrease in the NHEJ was observed when the catalytic domain of PRDM9 was fused to Cas9 (Supplementary Fig. 1A, gRNA-1 p-value=0.01 and gRNA-2 p-value=0.03). There was a significant increase of HDR/NHEJ ratio for the fusion with the HDR enhancer PRDM9 (Fig. 1E, p-value=0.007). Performance of PRDM9 fused with nuclease, nickase and dead variants of Cas9 was also tested. We observed negative results of nCas9-PRDM9 and dCas9-PRDM9 (Fig. 1F). The role of PRDM9 in enhancing the HDR outcome of the editing event was confirmed by fusing a catalytically dead variant of PRDM9 to Cas9 (Fig.1F). Absolute values are shown in Supplementary Fig. 1B.

### Synergistic HDR enhancer (SHE)

In order to explore additional factors that combined with PRDM9 could further increase the HDR efficiency we generated a Synergic HDR enhancer System or SHE (Fig. 2A) by employing a modified gRNA with a tetra-loop of phage sequence MS2 described elsewhere for synthetic transcriptional activation (Dahlman et al. 2015). This library consisted of multiple selected rad51 orthologues (Chintapalli et al. 2013) and rad52 (Paulsen et al. 2017) which are key in HDR, in addition to other HDR associated proteins such as CtIP, important in the resection of DNA after a DSB that have HDR enhancer activity when fussed to Cas9 (Tran et al. 2019), as well as other proteins such as adenovirus 4 proteins E1B-55K and E4orf6 which inhibit the competing pathway NHEJ (Chu et al. 2015). Other viral proteins like UL12, also stimulates recombination (Schumacher et al. 2012). Expression of a dominant mutant variant of 53BP1 (DN1S) also inhibit NHEJ, by suppressing 53BP1, a key regulator of DSB repair pathway choice in eukaryotic cells and functions to favor NHEJ over HDR (Jayavaradhan et al. 2019) (Canny et al. 2018). Dsup is a Tardigrade protein that modulates DNA damage repair and was also included (Ricci et al. 2021). DMC1 was also selected for the library, since this is a meiosis specific protein implicated in the repair of DSB using the homologous chromosome as the template for repairing the break (Hinch et al. 2019)(Cole et al. 2012). Members of this library are summarized in Supplementary Table 1.

**Figure 2.**
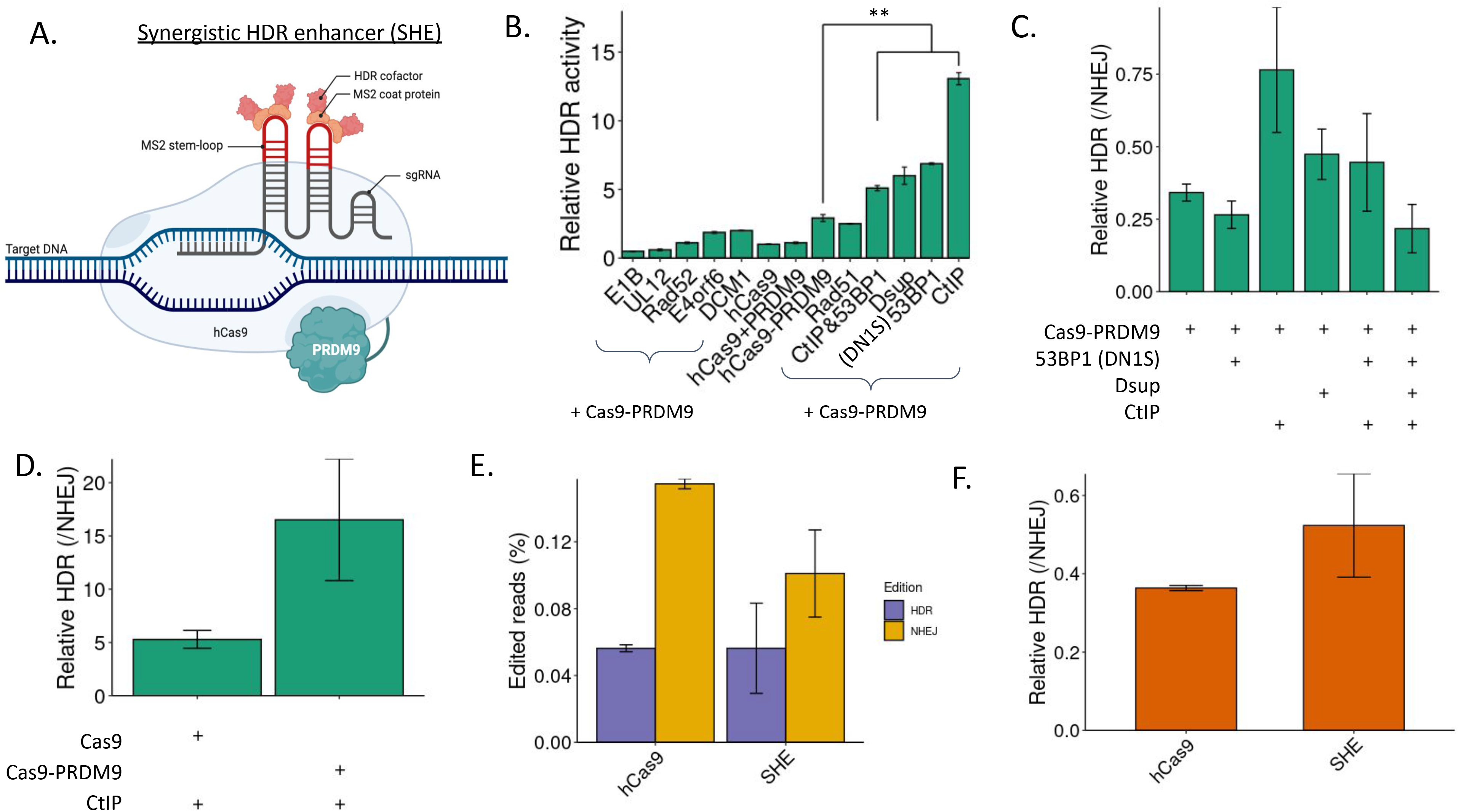
HDR enhancement by Cas9 fusion proteins. A) Scheme of synergistic HDR Enhancer technology: Cas9 (in blue) is fused to PRDM9 (green). Cas9 is associated with different HDR enhacers via sgRNA loops that include MS2 phage aptamer sequences bound by fusion proteins of HDR enhancers and the MS2 coat protein. B) HDR/NHEJ ratio of the Traffic Light Reporter, upon transfection with Cas9, Cas9-PRDM9, Cas9 co-deliver with PRDM9 and Cas9-PRDM9 together with HDR enhancing factors alone or in combination. C) HDR/NHEJ ratio of the Traffic Light Reporter, upon transfection with Cas9-PRDM9 and Cas9-PRDM9 together with the best HDR enhancing factors. D) HDR/NHEJ ratio of the Traffic Light Reporter, upon transfection with Cas9 or Cas9-PRDM9 together with MS2-CtlP. E) SHE performance for HDR in human iPS cells for HDR and NHEJ comparison, F) and relative measurements.

Split GFP reporter was used to compare the relative HDR activity of the different members of the library. As previously shown, there are significant differences between Cas9 and Cas9-PRDM9. Moreover, the co-delivery of Cas9-PRDM9 together with CtIP and 53BP1, Dsup, 53Bp1-DN1S or CtIP have a significantly improved HDR activity (Fig. 2B). Regarding the combination of 53BP1-DN1S, Dsup and CtIP with Cas9-PRDM9, higher levels of HDR are observed when adding CtIP, Dsup or both, but not in combination with 53BP1-DN1S when HDR/NHEJ ratio is determined with the traffic light reporter cell line (Fig. 2C). CtIP positive effect in combination with Cas9-PRDM9 was further confirmed in Fig. 2D, where we can see higher relative HDR in co-delivered with Cas9-PRDM9 than co-delivered with Cas9 alone.

The effect of Cas9-PRDM9 in relative HDR efficiencies has been also explored in human iPSC. Even though the results are not statistically significant, there is a decrease in NHEJ (Fig. 2E) that improves the HDR/NHEJ ratio (Fig. 2F).

### Template exploration with Cas9-PRDM9 using traffic light reporter

A library of multiple ssDNA donors was used to determine the best parameters in the donor design (Supplementary Fig. 2A). Left-asymmetric arm, with a left arm of 100 nucleotides and right arm of 50 nucleotides, and corresponding to the targeted strand, is the template that showed better HDR results after Cas9 activity (Supplementary Fig. 2B). After that, we checked if the results obtained with Cas9-PRDM9 correlated with those observed with only Cas9. For this, we choose a subsample of templates with the most promising donors representing the different groups. We used an asymmetric template with a left arm of 50 nts and right arm of 100 nts (t15), asymmetric right arm with same lengths (t27), and a symmetric template with 75 nts arms (t21). The sequence of three temples was the target strand, containing the PAM sequence. We also used the reverse complement sequence of template 27 (t28). Tendencies between Cas9 and Cas9-PRDM9 results with templestes representing the different groups of the library of templates are comparable (Supplementary Fig. 2C).

## DISCUSSION

Genome editing precision is mandatory when safe, controlled and precise modifications are required. Here we describe SHE, a synergistic HDR enhancer, that increases HDR precise outcomes due to the fusion of Cas9 with PRDM9 meiotic factor and co-delivery of CtIP that co-localizes with the gRNA though the MS2 coat protein linked to CtIP. We have tested the effectiveness of this system with small edits, as well as longer edits of multi-kilobase size. In both cases, we have achieved better results than using Cas9 alone. On one hand, we have concluded that PRDM9 fusion works better than the co-delivery of PRDM9 and Cas9, since the fusion facilitated the co-localization of both proteins in the target site. On the other hand, we have established that PRDM9 fusion with nickase Cas9 or dead Cas9 does not work, highlighting the essentiality of Cas9 nuclease activity, because double strand break is required for HDR to take place. PRDM9 with a catalytic mutation fused to Cas9 also showed a lower activity than wild type PRDM9 fusion, confirming the enhancing role in HDR of this factor.

The best performance has been achieved when Cas9-PRDM9 has been co-delivered with CtIP and colocalization with MS2 coat protein. Dsup also has shown a modest increase in relative HDR, while 53BP1-DN1S decreases the gain that the use of Cas9-PRDM9 implies. When all factors are transfected together, there is a loss of the synergistic effect, that could be due to unexpected interactions between the factors. The synergistic effect of CtIP together with Cas9-PRDM9 has been validated comparing its performance with Cas9. The synergistic effect of Cas9-PRDM9 and CtIP is 3-fold higher than Cas9 and CtIP. SHE has also been used in human iPSC, a relevant cell type for research and medicine.

In conclusion, in this study we have established an HDR enhancement system taking into account the combination of factors and optimal donor design. In the future, this system can be complemented with other HDR cofactors and be explored in different cell lines and organisms, accelerating biomedical research and the development of medicinal products to treat human diseases. The library of factors could also be extended by using comparative genomics in databases to further include other potentially important modulators of HDR.

## MATERIAL AND METHODS

### Cloning and plasmids

PRDM9 catalytic domain together with a flexible linker (GGGGS) was obtained from Twist Bioscience. PRDM9 was cloned into a Cas9-expression vector Addgene #41815 using isothermal assembly following standard protocols. Dead (D10A and H840A) and nickase (D10A) Cas9 mutations as well as PRDM9 catalytic mutation (G172A) were introduced by site directed mutagenesis (New England Biolabs). The collection of HDR enhancing factors fused to MS2 protein were constructed based on the pcDNA™3.1 vector backbone (Thermo Fisher). The following expression vectors were amplified by PCR: RAD51 (Addgene #41815), CtIP (Addgene #109403), Ad4orf2B (Addgene #64221), Ad4E4orf6 (Addgene #64221), 53BP1-DN1S (Addgene #131045), D-sup (Addgene #90019) and DMC1 (CRG ORFome collection) were amplified by PCR. Similarly, MS2 expressing vector (Addgene #61423) was also amplified by PCR. HDR enhancing factors and MS2 tag were cloned between Esp3I sites by Golden Gate Assembly (New England Biolabs). All the primers are listed in Supplementary Table 2. In the case of UL12, a gBlock of MS2-UL12 was synthesized by Twist Bioscience and cloned into pcDNA™3.1 vector backbone by Golden Gate Assembly. MS2-RAD52 was ordered as a plasmid vector to Twist Bioscience. gRNAs were cloned into sg-RNA(MS2) cloning vector (Addgene #61424) between BbsI sites using standard cloning methods. Sequences of gRNA TLR, gRNA AAVS1-3, previously described (Mali et al. 2013).

### Template library design

A library of 25 ssODN templates was designed to explore the effect of symmetry, strand, arm length and total template length (Supplementary Fig. 1A). Lengths of 25, 50, 75 and 100 nt were combined in both homologous arms that flanks the missing piece of the GFP to reconstitute the traffic light reporter (TLR). The templates with a free energy entropy lower than −33Gb were not synthetized since the predicted secondary structures were too strong to be used as a template with success. Regarding the total homological length, we explored lengths from 50 to 150 nt as a consequence of left and right arm length combinations.

### Cell culture, transfection and electroporation

Hek293T cells were infected with TLR lentivirus at a 0,2 multiplicity of infection (MOI) to generate the TLR cell line as described before (Certo et al., 2011). After infection cells were selected with puromycin following standard protocols for 2 weeks before being used for subsequent assays.

Hek293T cell line containing pT4 SMN1 2/2 emGFP was generated by PEI mediated transfection of SB100X and pT4 SMN1 2/2 emGFP DNA constructs, followed by single clone expansion and PCR genotyping. A positive clone was selected and expanded and used for subsequent assays. TLR cell line and C2C12 cell line (ATCC) were cultured at 37°C in a 5% CO 2 incubator with Dulbecco’s modified eagle medium (DMEM), supplemented with high glucose (Gibco, Therm Fisher), 10% Fetal Bovine Serum (FBS), 2 mM glutamine and 100 U penicillin/0.1 mg/mL streptomycin. The FiPSC-Ctrl1-Ep6F-5 cell line was purchased from the Biobank of the Barcelona Center for Regenerative Medicine (CMRB), and grown using standard conditions. Cell’s transfection experiments were performed with Polyethyleneimine (PEI, Thermo Fisher Scientific) at 1:3 DNA-PEI ratio in OptiMem. Cells were seeded the day before to achieve 70% confluence on transfection day (usually 120.000 cells in adherent p24 well plate). Plasmid molar ratio was 1 of Cas9 or Cas9-PRDM9: 3 gRNA: 3 HDR template using 0,089 pmols of Cas9 or Cas9-PRDM9 for a p24 well plate. For the experiments with the HDR enhancers plasmid molar ratio was 1 Cas9-PRDM9: 2 HDR enhancer: 3 gRNA: 3 HDR template using 0,089 pmols of Cas9-PRDM9 for a p24 well plate.

Electroporation of C2C12 and iPS cells was performed according to the manufacturer’s instructions with Lonza 4D-Nucleofector System with CD-137 program (C2C12). In the case of the iPS the parameters P3 Primary Cell 4D-Nucleofector were followed and the program CM-113 was applied.

### Analysis of HDR efficiency by FACS

When targeting HDR efficiency in TLR cells, targeted integration results in cells becoming GFP-positive which can be easily monitored by FACS analysis. Cells were analyzed by flow cytometry using BD LSR Fortessa (BD Bioscience. Blue 488nm laser with 530/30 filter and Yellow Green 561nm laser with 610/20 filter) 3-4 days after transfection. The relative HDR frequency was obtained by normalizing HDR by NHEJ frequency (RFP-positive cells).

### NGS Library Preparation

To quantify precise HDR genomic DNA was extracted using DNeasy Blood and tissue kit (Qiagen) and it was amplified using two-steps PCR and sequenced using Illumina sequencing platform (Miseq). DNA contains wild-type and edited DNA molecules, which were amplified together using the same pairs of primers (Supplementary Table 2). The two PCR reactions were performed with KAPA HiFi DNA Polymerase following manufacturer protocol: 100ng DNA, 0.3μM of forward and reverse primers in a final reaction volume of 25μl. Primers included sequencing adapters to their 3’-ends, adding them to both termini of PCR products that amplified genomic DNA. For the second PCR the primers used appended dual sample indexes and flow cell adapters. PCR products were purified with a PCR purification kit (Qiagen) and quantified using the QuBIT dsDNA High Sensitivity Assay kit and the QuBIT 2.0 fluorometer according to the manufacturer’s instructions (Life Technologies) before preparing the sequencing reaction.

### Bioinformatics analysis

Data coming from miSeq was analyzed using the CRISPR-A nextflow pipeline (Sanvicente-García et al. 2022). CRISPR-A template-based edits search is based on CIGAR as well as indels search. The quantification is done using the given template and the other counters are updated taking into account the nature of the change. First, the reference is modified using the template that should be shaped by the modification and homology arms in both sites of the change. Then, the new reference with the modification incorporated is aligned against the template with the same algorithm and parameters used to align the sample reads. Finally, reads are classified between wild type, indels or template-based in function of the alignment results against the reference amplicon and the modified amplicon.

### Statistical tests

Mean and standard deviation of replicates has been calculated using r-base function. Statistical significance has been determined though t-test when two groups are compared or ANOVA for more than two groups. Tukey Honest Significant Differences was used to determine which were the pairs of comparisons that have statistically significant differences.

## SUPPLEMENTARY FIGURE LEGENDS

Supplementary Figure 1. **HDR and NHEJ absolute measurements.** A) HDR and NHEJ absolute values measured by number of cells expressing GFP or RFP respectively after a double stranded break in the reporter cell line mediated by Cas9 or Cas9-PRDM9 fusion using two different gRNAs, or B) PRDM9 fused with nuclease, nickase and dead variants of Cas9 as well as catalytic mutant PRDM9.

Supplementary Figure 2. **Template exploration with traffic light reporter system for HDR and NHEJ.** A) Scheme of ssDN library members. Dot lines goes from the modification position, straight bold line, and are separated by 25nts each. B) HDR/NHEJ ratio of the Traffic Light Reporter using template library upon Cas9 activity. B) Comparison between Cas9 and Cas9-PRDM9 using different templates representatives from the design parameters. C) A subset of 4 templates is used to compare Cas9 with Cas9-PRDM9. We used an asymmetric template with a left arm of 50 nts and right arm of 100 nts (t15), asymmetric right arm with same lengths (t27), and a symmetric template with 75 nts arms (t21). The sequence of three temples was the target strand, containing the PAM sequence. We also used the reverse complement sequence of template 27 (t28).

## Authors contributions

MG and ASM conceived the project and designed the experiments. LGB performed in vitro experiments. LB and SC designed and performed iPSC experiments. MSG performed genome editing analysis, and designed ssDNA donor library. ASM and MSG, wrote the initial version manuscript. All authors discussed the results and contributed to the manuscript writing process

## DATA AND MATERIALS AVAILABILITY

Next-generation sequencing data are available in the European Nucleotide Archive under the Study accession number PRJEB58704. Custom analysis scripts for data analysis and visualization are freely available at https://bitbucket.org/synbiolab/prdm9_figures/.

