## Supplementary figures and images for "Synergic homology directed recombination by PRDM9 meiotic factor"

### Supplementary Figures

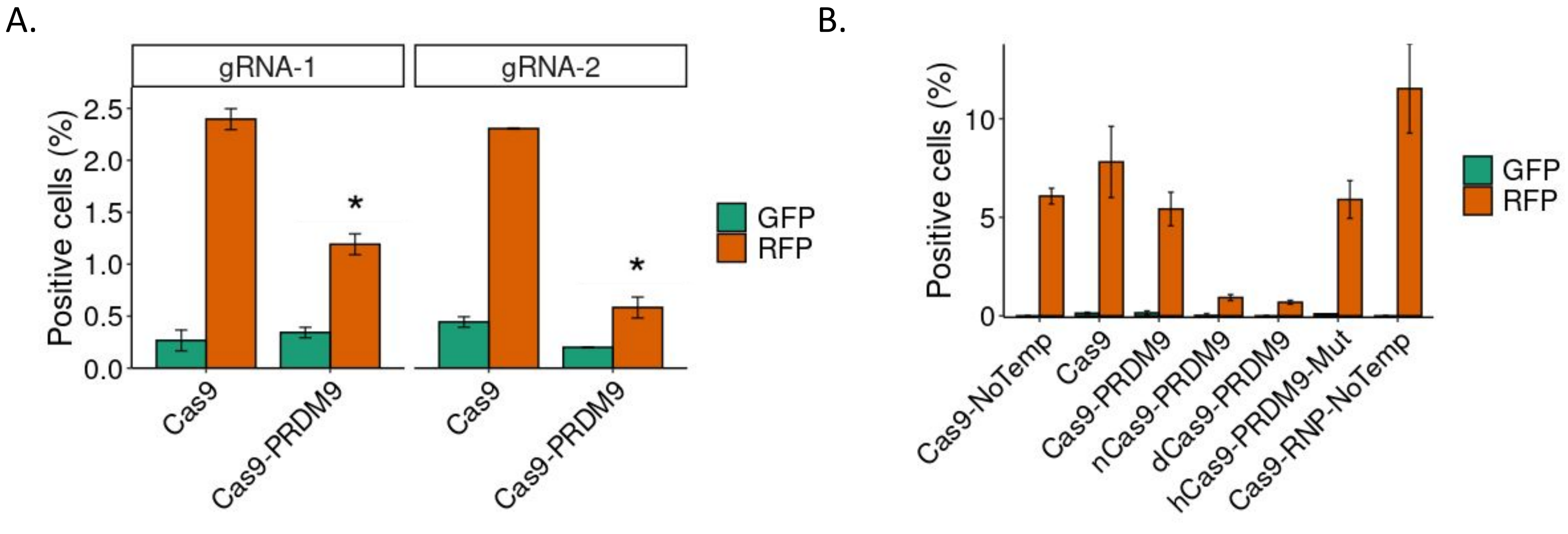

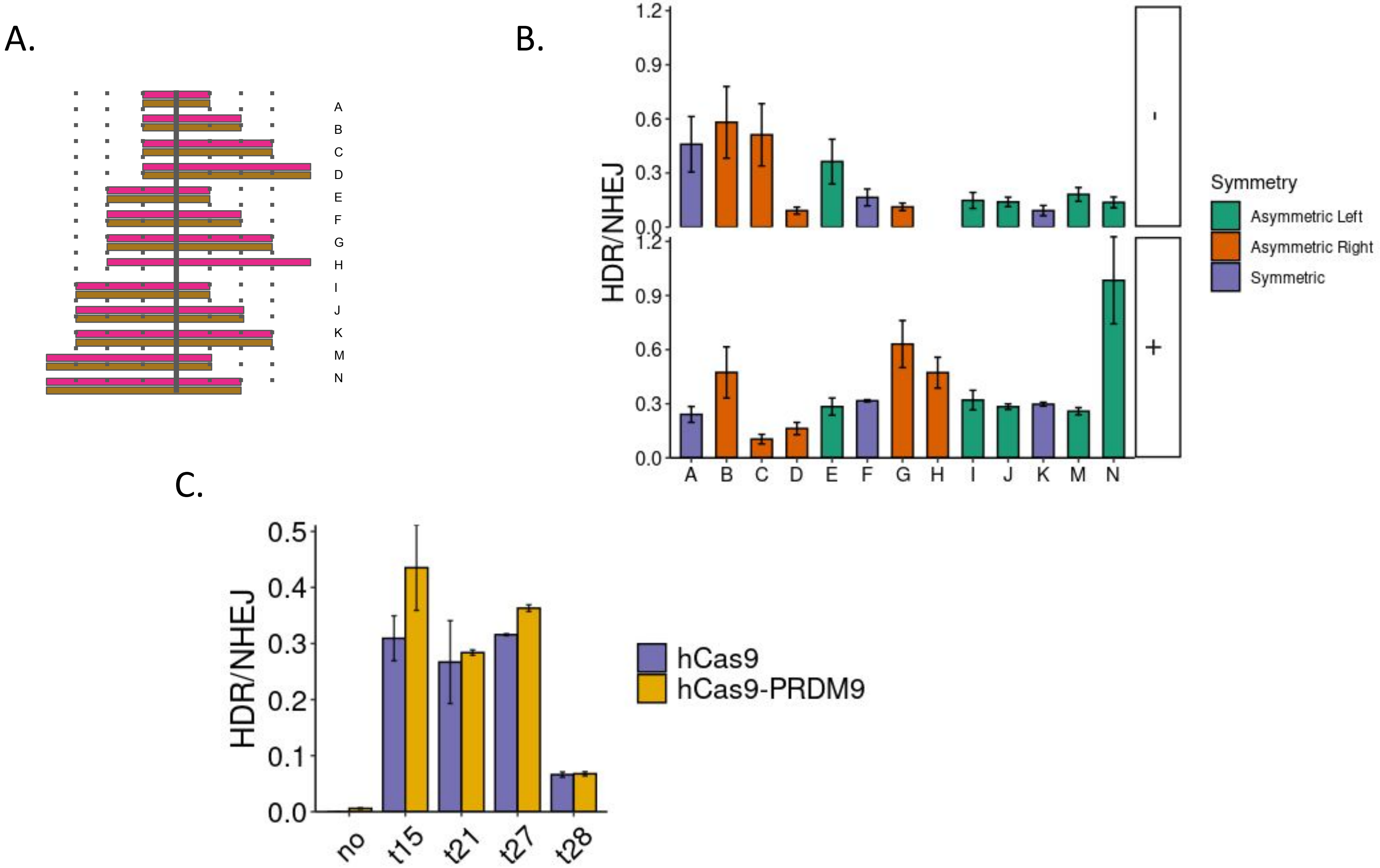
